# Ancestral Sequence Reconstruction as a tool to detect and study de novo gene emergence

**DOI:** 10.1101/2024.01.02.573862

**Authors:** Nikolaos Vakirlis, Omer Acar, Vijay Cherupally, Anne-Ruxandra Carvunis

**Affiliations:** Institute for Fundamental Biomedical Research, BSRC “Alexander Fleming”, Vari, Greece; Pittsburgh Center for Evolutionary Biology and Medicine, Department of Computational and Systems Biology, School of Medicine, University of Pittsburgh, Pittsburgh, PA 15213, United States

## Abstract

New protein-coding genes can evolve from previously non-coding genomic regions through a process known as de novo gene emergence. Evidence suggests that this process has likely occurred throughout evolution and across the tree of life. Yet, confidently identifying de novo emerged genes remains challenging. Ancestral Sequence Reconstruction (ASR) is a promising approach for inferring whether a gene has emerged de novo or not, as it can enable us to inspect whether a given genomic locus ancestrally harbored protein-coding capacity. However, the use of ASR in the context of de novo emergence is still in its infancy and its capabilities, limitations, and overall potential are largely unknown. Notably, it is difficult to formally evaluate the protein-coding capacity of ancestral sequences, particularly when new gene candidates are short. How well-suited is ASR as a tool for the detection and study of de novo genes? Here, we address this question by designing an ASR workflow incorporating different tools and sets of parameters and by introducing a formal criterion that allows to estimate, within a desired level of confidence, when protein-coding capacity originated at a particular locus. Applying this workflow on ∼2,600 short, annotated budding yeast genes (<1,000 nucleotides), we found that ASR robustly predicts an ancient origin for most widely conserved genes, which constitute “easy” cases. For less robust cases, we calculated a randomization-based empirical P-value estimating whether the observed conservation between the extant and ancestral reading frame could be attributed to chance. This formal criterion allowed us to pinpoint a branch of origin for most of the less robust cases, identifying 33 genes that can unequivocally be considered de novo originated since the split of the *Saccharomyces* genus, including 20 *S. cerevisiae*-specific genes. We find that the remaining, equivocal cases, may be explained by different evolutionary scenarios including rapid evolution and multiple losses, as well as a very recent de novo origin. Overall, our findings suggest that ASR is a valuable tool to study de novo gene emergence but should be applied with caution and awareness of its limitations.

## Introduction

How new genes originate is a fundamental question in biology because genetic novelty underlies molecular, phenotypic and organismal novelty^1^. Understanding how and when novel genes arise is therefore essential to understand evolution at every level of biological organization. For a long time, new genes and protein functions were believed to result exclusively through tinkering and recombination, using pre-existing genes and gene parts as raw material^2^. Consequently, processes such as substitutions, duplication and divergence, gene fusion and fission, exon shuffling, or horizontal gene transfer (HGT) have been extensively studied and their importance is established.

Nonetheless, a radically different route to genetic novelty exists: a novel gene can evolve from entirely non-coding sequences in a process known as de novo gene emergence^3,4^. Long considered so improbable as to be impossible^5^, de novo gene emergence has high potential to produce an entirely new protein function, since the emerging protein will be free of constraints present in pre-existing, already functional sequences. De novo genes have been found in every eukaryotic lineage studied so far and can have central, even essential cellular functions^4^. This has led to de novo emergence being increasingly viewed as a universal evolutionary mechanism.

It is challenging to distinguish whether a new gene has emerged de novo or through other evolutionary processes. Indeed, rapid sequence divergence beyond recognition following events such as duplications or rearrangements, as well as HGT can also result in a gene appearing to be novel or taxonomically restricted^6,7^. It is thus important to develop robust methods for distinguishing between these different evolutionary routes, in order to be able to assess the impact of de novo gene emergence and study the characteristics and function of de novo genes.

Evidence for de novo gene emergence can be provided by computational comparative genetics approaches^8^. The strongest kind of evidence is confidently inferring that the genomic locus that now harbors a novel gene in a given lineage, did not encode a protein sequence in the past. Until recently, the gold standard was to align the novel gene to its orthologous regions in multiple outgroup species and demonstrate that these outgroup loci were noncoding by identifying specific mutations that enabled the presence of an open reading frame (ORF) in the genome of interest. Parsimoniously one can then infer that the ancestral state of the positions in question was such that disrupted the ORF currently present in the focal lineage.

However, visually inspecting such alignments does not always result in clear cut conclusions. It is especially delicate to draw robust conclusion when the candidate new genes are short and when alignments contain frameshifts, and in such tricky cases conclusions are typically drawn based on a personal judgement call rather than a formal test. A potentially more powerful and accurate approach is to use Ancestral Sequence Reconstruction (ASR). ASR allows both estimating whether an ancestral sequence contained an ORF or not as well as tracing the mutational transition from noncoding to coding. Thus, the application of ASR to the task of de novo gene detection could be a potent tool to gain insights into this evolutionary process.

ASR has mostly been used in the context of protein alignments in order to reconstruct ancestral protein sequences from extant ones^9^. Only a handful of studies have applied ASR to de novo genes^10–16^ thus far, and an in depth assessment of its performance and limitations is lacking. Here, we evaluate how reliably existing ASR tools can estimate the emergence date of short ORFs annotated in the budding yeast *S. cerevisiae*’s genome. Using reading frame conservation (RFC) between an ORF and its inferred ancestor as a quantitative measure of ORF age, we conclude that ASR allows robust evolutionary inference for ancient genes, but shall be used with caution to infer recent events of de novo gene emergence.

## Results

### A computational pipeline to reconstruct and conservatively estimate the coding capacity of ancestral nucleotide sequences

We assembled nucleotide multiple sequence alignments of all annotated *S. cerevisiae* ORFs shorter than 1000nt (n=2,640, Figure 1A) with their respective orthologous genomic loci in closely related Saccharomyces species (see Methods and Figure 1B). These alignments served as inputs for ASR using a pipeline we designed to test how robust ASR inferences would be to methodological choices. This pipeline combines three different phylogenetic tools, phylogenetic trees built with and without the species topology as a constraint, and two different types of input alignments: one based only on the exact region of the *S. cerevisiae* ORF and one based on an extended region (see Methods). The phylogenetic tools are: FastML^17^ which performs both marginal and joint Maximum Likelihood (ML) reconstructions of characters and ML reconstruction of indels (hereafter FastML_joint and FastML_marginal), PREQUEL from the PHAST^18^ package, which performs ML reconstruction of characters and parsimonious reconstruction of indels and prefers to infer deletions when insertions and deletions cannot be distinguished (producing an upward bias on the length of the sequence at root) and PRANK^19^ which uses ML reconstruction of characters and a custom algorithm for the inference of insertions and deletions (Figure 1A).

**Figure 1.**
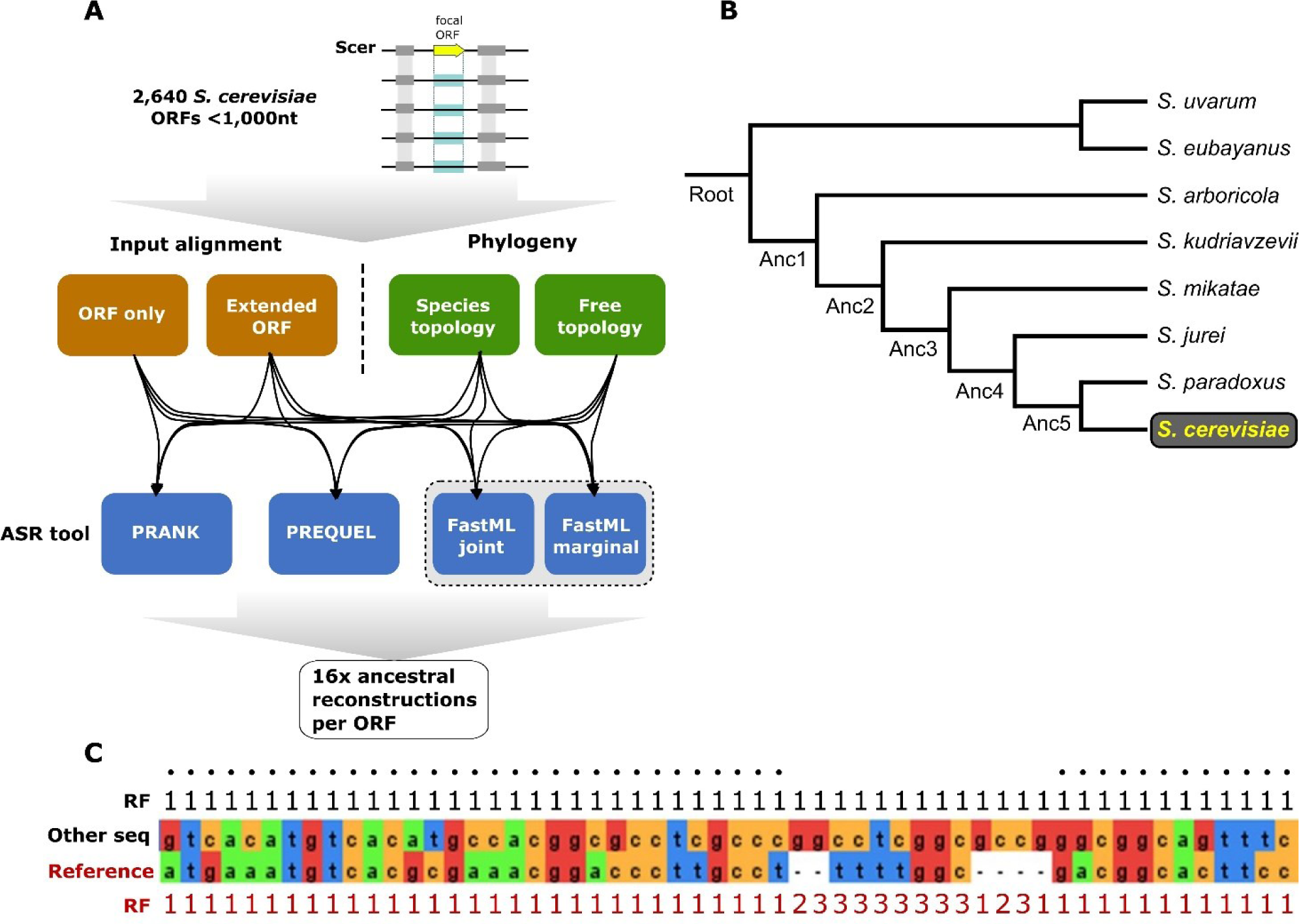
**A:** Overview of the workflow followed in the present study to generate a set of ASR methodological variations. **B:** Tree topology of the Saccharomyces genus used in our analyses. **C:** An example alignment to illustrate how the Reading Frame Conservation measure is calculated. The reading frame (RF) of each sequence and how it changes with the presence of gaps, is shown. RFC is calculated by counting non-gap positions where the two frames match (marked with blacked dots) and dividing by the length of the reference sequence, which in this study is always the *S. cerevisiae* ORF.

After running our ASR pipeline on all input alignments, we searched each ancestral sequence for the presence of ORFs that could correspond to ancestral versions of the extant *S. cerevisiae* gene. We defined these putative ancestral ORFs in two ways: ATG-STOP hereafter “ATG”, or STOP-STOP hereafter “noATG”. The similarity of each ancestral ORF longer than 30nt relative to the extant one of *S. cerevisiae* was scored using the Reading Frame Conservation (RFC) measure^20,21^: (length covered by the ancestral ORF aligned in the *S. cerevisiae* ORF frame)/(length of the S. cerevisiae ORF). An RFC value = 1 means that an ancestral ORF exists that is at least as long as the *S. cerevisiae* ORF and aligns to it in the same frame and without frameshifts (see Figure 1C for an example). We inferred whether a *S. cerevisiae* ORF originated de novo since the Saccharomyces common ancestor according to each combination of methodological choices implemented in our pipeline, for a range of RFC cutoffs (0.5, 0.6, 0.7, 0.8).

We compared the outputs of ASR with a classification of the input ORFs into emerging or established previously developed by Vakirlis et al. using a combination of sequence and selection signatures^11^. Figure 2 shows the results when using the definition of ORF without the need for an ATG start codon (noATG). Results using the alternative definition are highly similar and can be found in Supp. Figure 1.

**Figure 2.**
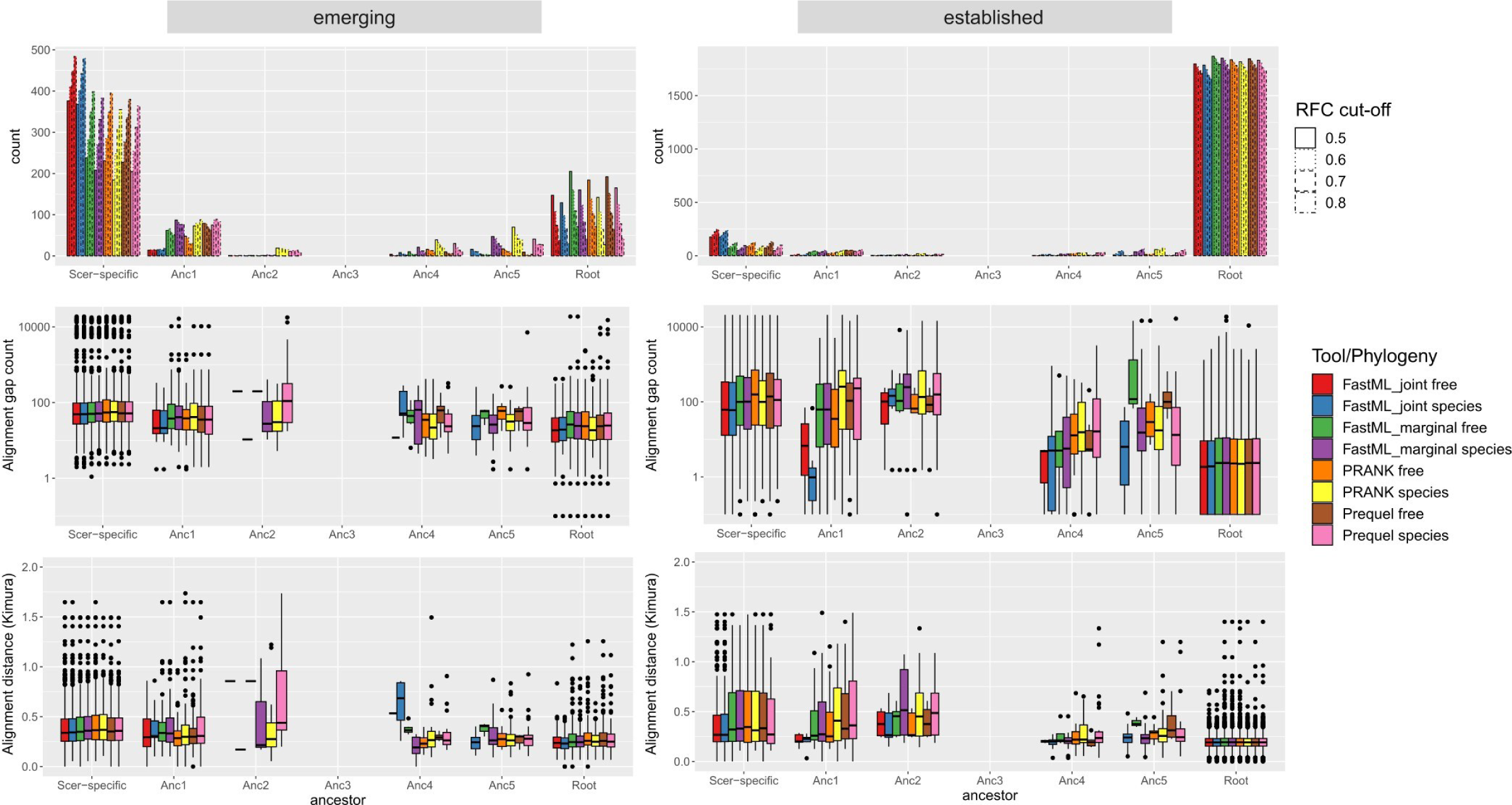
Distributions of branches of origin and input multiple sequence alignment statistics. **Top**: Distributions of branch where the most ancient ancestral ORF has been identified using four different RFC cut-offs and eight different ASR methodologies (tools+phylogeny). Bars within each node (e.g. “Root”) correspond, from left to right, to RFC cut-off of 0.5, 0.6, 0.7 and 0.8. Results for ORF-only alignments and “noATG” definition of ORFs are shown. Results for extended ORF alignments and ATG can be found in Supp. Figure 1. **Middle**: Distributions of average gap count per sequence in the initial multiple sequence alignment, over the different predicted branches of origin (most ancient ancestral ORF, using an RFC cut-off of 0.5). **Bottom**: Same as above, but for the average pair-wise Kimura distance in the input alignment.

In agreement with the initial analysis, the vast majority of established ORFs were classified by ASR as ancient and the majority of emerging ORFs were classified by ASR as *S. cerevisiae* - specific. The type of initial alignment used as input (ORF only or extended ORF region) had minimal impact, with the vast majority of ORFs being predicted to have the same origin. When setting an arbitrary RFC cut-off of 0.5, ORFs predicted to be ancient by ASR have initial alignments with shorter genetic distance and more gaps, than those predicted to be *S. cerevisiae* - specific (Figure 2). This is consistent with young ORFs evolving faster, which has been reported before^22–24^, but it can also be explained with faster evolving ancient genes whose more “difficult” initial alignments are challenging for ASR.

ASR methodologies unequivocally inferred 1,355 ORFs to be at least as ancient as the Saccharomyces ancestor, with RFC>0.9. Almost all these ORFs (1351/1355) were also classified as “established” by Vakirlis et al. Overall, these 1,355 cases can safely be considered ancient and we conclude that ASR performs well in such cases (Supp. Table 1).

### An empirical P-value allows to confidently select a most likely branch of origin for many ORFs

For the remaining 1,275 ORFs, different ASR methodological variations gave at least partly conflicting estimates. Two examples of such ORFs and the best RFC scores in ancestral sequences predicted by different methodologies can be found in Figure 3A. On the left we show a case where, using an RFC cut-off of 0.5, some methodologies would predict a much more ancient origin than others. On the right, we show a case where no methodology retrieves an ORF with RFC>0.4 in any ancestor, thus using an RFC cut-off of 0.5 would result in a coherent classification as species-specific, across ASR methodological variations, but an RFC cut-off of 0.3 would not. Relying on an arbitrary RFC cut-off for inferring ORF origination is problematic and fickle.

**Figure 3.**
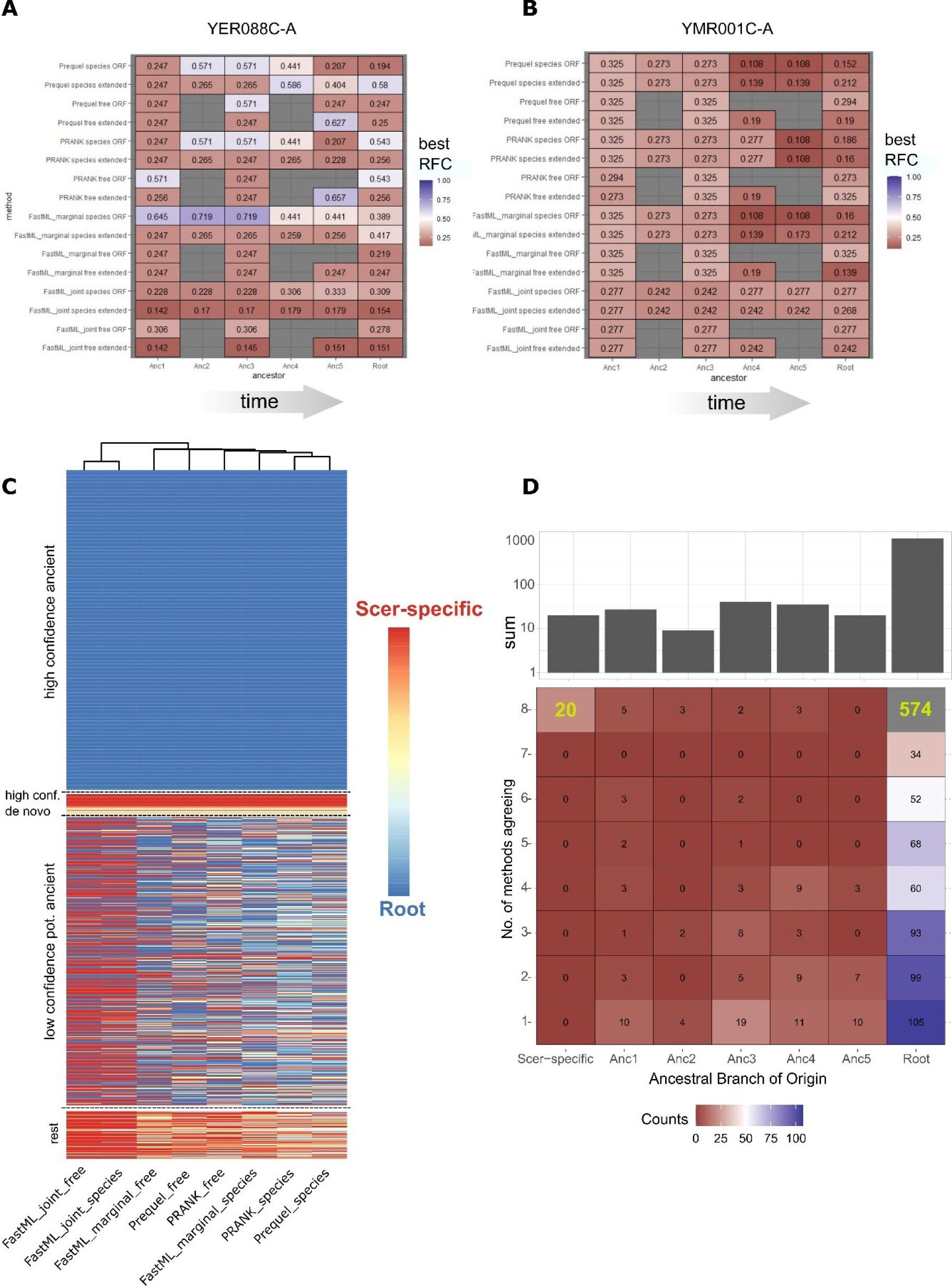
**A:** One example of maximum RFC score for different ancestors, as predicted using ancestral sequences from different methodologies (tool+phylogeny+inp. alignment). noATG ORF definition shown. Example is gene YER088C-A, a representative case where picking a branch of origin is not immediately obvious since when using an arbitrary RFC cut-off (e.g. 0.6) some methodologies lead to presence of ORFs in much older nodes than others; for example FastML_marginal+species+ORF would place the origin at Anc3, whereas PRANK+species+extORF would lead to a species-specific origin. **B**: Another example (YMR001C-A) where no methodology predicts presence of an ancestral ORF, even with an RFC cut-off as low as 0.5, and should thus be considered species-specific. **C**: Heatmap showing the ancestral branches of origin as defined using our empirical P-value cut-off of 0.01, for each of the 1,275 ORFs that were not previously predicted to be robustly ancient, using 8 different ASR methodologies (ORF-only alignments, noATG ORF definition). Based on the agreement among methodologies, we can group them into high-confidence ancient (top group), high-confidence *de novo* originated, including *S. cerevisiae*-specific (second from the top), low-confidence but potentially ancient since at least one methodology predicts and ancient origin (third from the top) and the rest (bottom). **D**: Number of methodologies agreeing in the ancestral branch of origin of each ORF. Same data as in C.

We thus asked whether one can systematically infer when an ORF has formed, taking into account the fact that ORFs can form randomly. To this end, we randomized each entire ancestral sequence while keeping its nucleotide composition constant and then logged the best RFC-scoring ORF, repeating this procedure 1000 times (see Methods). This produced an empirical distribution of values which we then used to assign a P-value on the best RFC-scoring ORF of the real ancestral sequence. The most ancient phylogenetic node where a P-value <0.01 was recorded was then kept as the most likely branch of origination of this ORF, that is, the most ancient branch where the presence of such an ORF is unlikely to be due to chance. It is important to stress that, ultimately, this approach may lead to strongly conservative estimates because de novo emergence could start from genomic loci which do happen to harbor unusually long ORFs, something that is bound to occur given that the genomic space is continuously explored throughout evolutionary time. The relationship between the empirical P-value and the best RFC ORFs at the root ancestor, which appears to have a sigmoid-like shape, can be found in Supp. Figure 2.

An overview of the predictions of branches of origin of the different methodologies when relying on the empirical P-value can be found in Figure 3C and the raw data can be found in Supp. Table 2. Note that since results between the ORF-only and extended-ORF alignments were highly similar, we only use the ORF-only alignments for this and all downstream analyses. When comparing the predictions of the different methodologies, FastML_joint stands out as resulting in more species-specific estimates than the rest (380 ORFs or 30% compared to 162 ORFs or 12.7% on average for the rest). This is because this method has a strong tendency to infer deletions when encountering gaps in the alignment, resulting in longer ancestral sequences. No other significant bias among methodologies was found.

We observed that for 574 ORFs, an ancient origin was predicted by all ASR methodologies, and we classified these as “High-Confidence” (HC) and ancient (Figure 3C, top group). For another 33 HC ORFs, all methodologies agreed on an origin after the split of the genus and so we can safely conclude that these have emerged de novo (Figure 3C; HC- de novo). Notably, 20 of them are *S. cerevisiae*-specific. Integrating ASR with a systematic RFC p-value therefore improved the robustness of evolutionary inferences relative to an arbitrary RFC cutoff. However, the remaining 667 ORFs should be considered “Low-Confidence” (LC) with discordant predictions that suggest uncertainty about when they originated, including 508 where at least one methodology predicted an origin at the root. In Figure 3D we provide a condensed view of these findings, allowing to compare the numbers of confident predictions, as well as those of more uncertain ones. We next asked what could explain this uncertainty.

### Multiple evolutionary scenarios could account for the uncertain origin of low-confidence ORFs

We compared the properties of HC-ancient ORFs, where all methodologies agreed that the node of origin is the node of the tree (n=574), with those of LC – potentially ancient ORFs where at least one, but less than four methodologies predict an origin at the root (n=297).We found that HC ancient ORFs are on average longer and have more similar initial alignments which also contain fewer gaps than LC ancient ones (Figure 4A). This supports the status of the high-confidence ones as more robustly ancient since they are suggestive of conservation of a true protein-coding ORF. LC ORFs have also significantly lower maximum RFC-scores in their root reconstructed ancestor (Figure 4A). In other words, either their best reconstructed ORFs are much shorter than the extant *S. cerevisiae* one, or they are long but don’t align well on the same frame as the extant *S. cerevisiae* ORF. Note here that 70 of the HC ORFs have relatively low best RFC score (<0.5), yet thanks to our empirical P-value we are able to confidently classify them as ancient. Finally, the posterior probabilities (as predicted by FastML_marginal) of the root reconstruction are lower both for indels and for individual positions (Figure 4A). Thus, LC ORFs should be viewed as much harder cases than HC ones, with borderline predictions of their phylogenetic origins.

**Figure 4.**
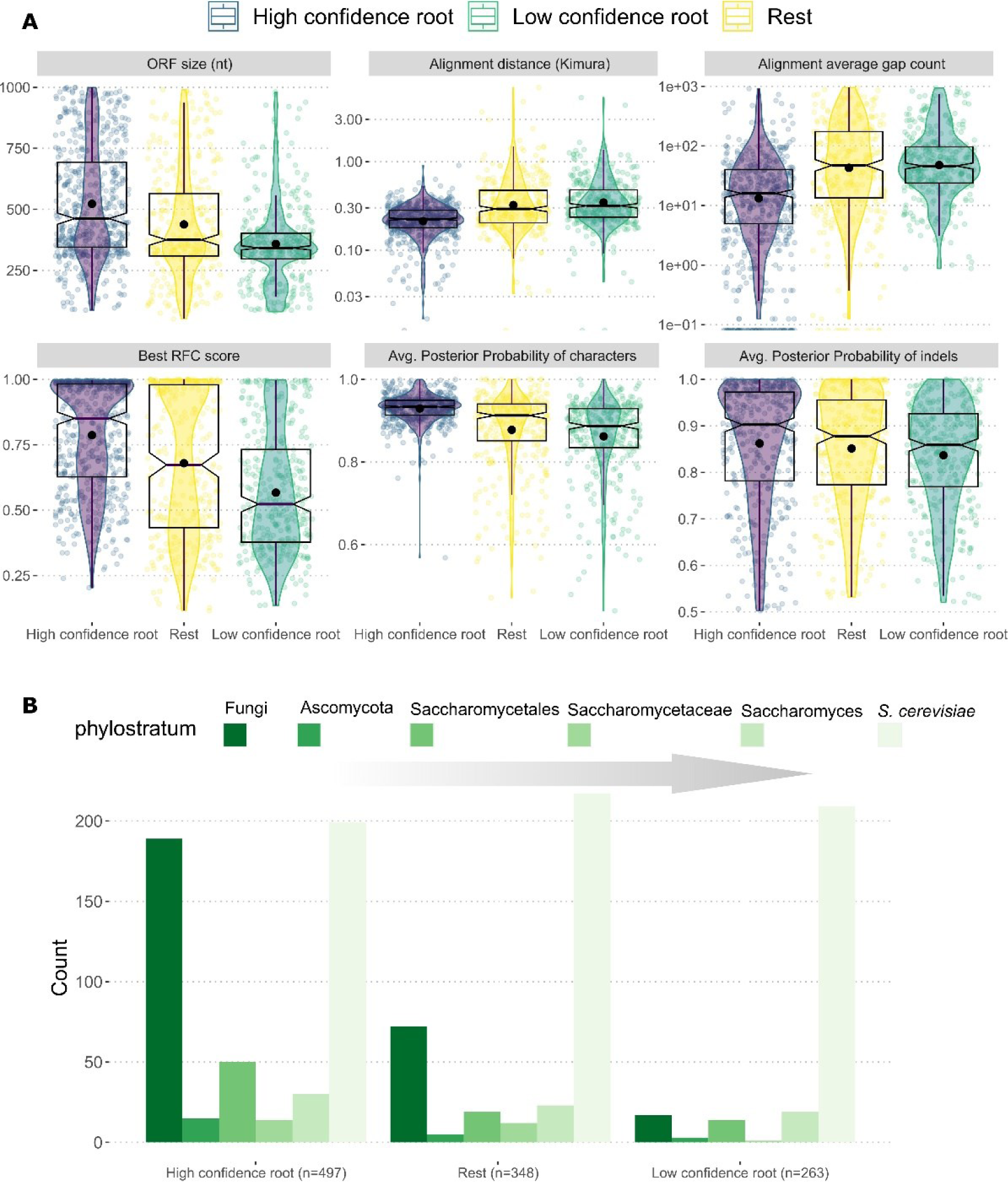
Comparison of low-confidence and high-confidence ancient ORFs. **A**: ORF size, best RFC score at root ancestor, number of gaps per sequence in the initial ORF-only MSA, average pairwise Kimura distance in the initial ORF-only MSA, average posterior-probability of characters at the root ancestor as calculated by FastML_marginal, average posterior probability of indels as calculated by FastML_marginal. **B:** Counts of ORFs per major phylostratigraphic origin for HC ancient ORFs, LC ancient ORFs and the rest. Bars are ordered from ancient to recent, left to right. For visual purposes, only phylogenetic branches corresponding to named taxonomic groups are shown.

While informative regarding the limitations of ASR, these differences between HC and LC ancient ORFs does not allow to propose a most likely evolutionary scenario for the origin of the LC ORFs. Indeed, we would expect to see the same set of differences if LC ORFs were fast evolving ancient ORFs for which only a minority of methods happen to succeed in capturing their ancient status, or if they were recently emerged ORFs from regions of the genome where the probability of forming an ORF was high, making detection of a long enough in frame ORF in one of the reconstructions more likely.

A relatively independent approach to estimating the timing of origination of a gene is protein-level phylostratigraphy^25^. We analyzed protein-level sequence similarity searches against all available fungal proteomes (see Methods) and obtained, for each *S. cerevisiae* ORF, a phylostratigraphic node of origin (most recent common ancestor of species with match in the fungal tree). In Figure 4B, we show the distribution of phylostratigraphic origins in the HC and LC ancient ORFs as well as for the other ORFs analyzed.

Most of the LC ORFs were *S. cerevisiae*-specific according to phylostratigraphy (79%). This percentage was significantly lower for HC (40%), as expected (Wilcoxon test P-value<2.2*10^-16^). However for these HC ORFs our ASR based analyses point to a phylogenetic origin that is strikingly older than the one recovered by phylostratigraphy. By contrast, we find a much higher percentage among the HC that have an ancient phylostratigraphic origin (38% at Fungi) compared to the LC ones (6.4%, Wilcoxon test P-value<2.2*10^-16^). These trends suggest that the HC and LC groups are qualitatively different while providing a validation of the ASR approach. Yet the origins of the LC group remain equivocal, because these results are what we would expect if they were mostly ancient but fast evolving, perhaps also easily lost genes, but it is also what we would expect if most of them were of truly recent origin.

## Discussion

ASR is a promising approach allowing to peek into the evolutionary past of sequences and elucidate the process of de novo gene origination. It has the potential to provide important novel insights both into the frequency and the evolutionary forces that drive de novo gene emergence. Nonetheless, it has been demonstrated that ASR is sensitive to many factors, including the methodology of multiple sequence alignment and the phylogeny^26,27^. In addition other biases might come in play when the ancestral sequences are examined for the presence of relevant ORFs in the context of de novo emergence. For these reasons, we performed a systematic examination and assessment of ASR for the study of de novo gene origination.

Overall, we find that ASR is well suited to be used as a tool for de novo inference and that the variability in the results from the different methodologies is limited. With the notable exception of the marginal reconstructions of FastML, the rest of the methodologies were for the most part in good agreement as per the node of origin of an ORF relative to random expectations. It is possible that, while in the context of ancestral protein sequence resurrection slight variations might lead to changes in functionally critical amino acids (e.g. that could affect the active site of a resurrected enzyme), the same slight variations might not impact the inference of the timing of origination of an ORF in the context of the present study.

An important point is that while we had one part of a good positive control set in genes with widespread presence and protein-level conservation in other species, we lack an independently generated “gold standard” set of de novo genes to compare to, but most importantly we lack an appropriate negative control. A potential solution to the latter would be to establish a gold standard collection of pseudogenes, or if this is not available, generate them through evolutionary simulations. These would represent evolution in the opposite direction of de novo emergence (gene death vs. gene birth) and they would be valuable as a test to our ASR-based workflow. For what percentage of such pseudogenes would we be able to accurately reconstruct their ancestral protein-coding status and the timing of their pseudogenization? This could be the focus of future work. At the same time, such work could also address the generalizability of the present findings in other lineages, including ones experiencing slower or faster evolution.

We believe that it is always best to start from a conservative place. Our empirical P-value approach might in fact be *too* conservative in considering all in frame ORFs longer than would be expected randomly as potential evidence for selection. This is inextricably linked to the poorly understood questions of how de novo gene emergence begins, at which point during de novo gene evolution a protein is first expressed, and at which point the incipient ORF is subjected to selection at the level of its size. If the initial evolutionary “version” of a de novo gene has on average the same length as any spurious small ORF on the genome then our assumption and the empirical P-value approach would be valid. But if a slightly longer than usual small ORF is mostly what de novo emergence starts from, then it would be strongly conservative, since such ORFs continuously appear and disappear in the genome throughout evolution. One can envision a future approach that incorporates this probability into the calculation of a P-value, to perhaps make it more realistic.

## Materials and Methods

### Description of the dataset

Our initial dataset consisted of 2,816 annotated protein-coding genes of *S. cerevisiae* that were included in the dataset analyzed by Vakirlis et al.^11^ and were under 1000nt long. The orthologous regions for each of these genes in seven Saccharomyces species (*S. paradoxus*, *S. mikatae*, *S. kudriavzevii*, *S. uvarum*, *S. jurei*, *S. arboricola*, *S. eubayanus*) were identified as follows: First, genomes were obtained from the following sources: *S. paradoxus* from ^28^*, S. arboricolus* from ^29^, *S. jurei* from ^30^ and *S. mikatae*, *S. bayanus var. uvarum*, *S. eubayanus*, and *S. kudriavzevii* from ^31^. Alignments were constructed between each *S. cerevisiae* ORF and its homologs in each Saccharomyces relative using synteny information. To identify anchor genes for syntenic blocks, BLASTP^32^ was run for each annotated ORF in *S. cerevisiae* against each ORF in the comparison species. Identified homolog pairs with E-value < 10^-7^ were selected as potential anchors. For each ORF in the *S. cerevisiae* genome, the upstream anchor G0 and downstream anchor G1 were selected that minimized the sum of the distance between the anchors in S. cerevisiae and the distance between the anchors in the comparison species; this sum was required to be less than 60 kb. The sequence between and including G0 and G1 were then extracted from both the *S. cerevisiae* genome and the comparison species and a pairwise alignment of the syntenic region was generated using MUSCLE v. 3.8.31. Multiple sequence alignments of the exact *S. cereivisiae* gene locus to its orthologous Saccharomyces genomic regions were generated with MAFFT^33^ using default parameters. We removed 176 *S. cerevisiae* genes from the dataset because the orthologous region of *S. cerevisiae* could only be identified in less than 6 species, or it contained >1000 gaps/sequence. For the remaining 2,640 genes, we also generated an alternative, extended version of the multiple sequence alignments by including 500nt flanking the *S. cerevisiae* ORF downstream and upstream. For each *S. cerevisiae* gene, we collected gene and protein properties from Carvunis et al^34^. and Vakirlis et al^11^. We also performed protein-level sequence similarity searches for every gene, against a protein sequence database containing all fungal proteomes downloaded from NCBI’s RefSeq in May 2021 plus the 332 *Saccharomycotina* proteomes from Shen et al.^35^. This search was conducted with BLASTp^32^ using an E-value cut-off of 0.001 and the *-max_target_seqs* flag set to 1000. The results of the similarity searches were processed as in^36^: For each gene, we first obtained the list of all fungal species with a significant similarity match. Phylogenetic age of each gene was then calculated as the most recent common ancestor of all species with a match. The NCBI Taxonomy common tree was used for this, resulting in classification into the following phylogenetic ages: species-specific, genus (*Saccharomyces*), family (*saccharomycetaceae*), order (*saccharomycetales*), division (*ascomycota*) or kingdom (*fungi*). For each gene, we also counted the number of species with match (number of species with homologues).

### Phylogenetic reconstruction

Phylogenetic trees were reconstructed using RAxML next generation^37^ (*raxml-ng*) with the GTR substitution matrix, empirically estimated rates and nucleotide frequencies, 4 categories of rates drawn from a GAMMA distribution with maximum likelihood inference of its shape parameter using the following command: *raxml-ng --seed 12546582 --model GTR+F+G*. For the species-topology phylogeny we additionally used the species topology as shown in Figure 1B in newick format with the additional arguments: *--evaluate –tree SACCH_TOP.nwk*. The resulting phylogenetic tree, in the case of the species-topology, was re-rooted using *S. eubayanus* and *S. uvarum* (Seub-Suva) as outgroups with the GoTree^38^ utility to ensure downstream consistency. The free topology tree was re-rooted at midpoint using GoTree. For use with the ancestral sequence reconstruction tool PREQUEL (see next subsection), a phylogenetic model had to be generated using the PHAST^18^ utility PhyloFit. The trees generated by RAxML were provided to PhyloFit together with the initial input MSA. We then confirmed that the tree and model fitted with PhyloFit was identical to that of RAxML.

### Ancestral Sequence Reconstruction

ASR was performed with FastML^17^ using the following command*: perl FastML_Wrapper.pl --MSA_File "INPUT_ALIGNMENT.fasta" --seqType nuc --Tree "INPUT_TREE.nwk" --SubMatrix GTR --OptimizeBL no --indelReconstruction ML –outDir “OUTPUT_DIR”*

ASR was performed with PRANK^19^ without iterations using the following command: *prank -d "INPUT_ALIGNMENT.fasta" -support -showall -keep -F -once -o="OUTPUT_PREFIX" -t="INPUT_TREE.nwk"*

ASR was performed PREQUEL from the PHAST^18^ package using the following command*: prequel " INPUT_ALIGNMENT.fasta" "INPUT_MODEL.phylofit_corTree.mod" OUTPUT_PREFIX,* and then once again with the *-n* argument to obtain the posterior probabilities.

### Identification of ORFs in ancestral sequences

For every ancestral sequence in each variation of ASR, we performed the following: first, we identified all ORFs on the forward strand using *getorf* from EMBOSS^39^ defined either as ATG-STOP or STOP-STOP. The coordinates of each ORF on the ancestral sequence were stored. Then, a pairwise alignment of the entire ancestral sequence and the *S. cerevisiae* extant ORF was generated using the command *“pairwise2.align.globalds(S.cer_sequence, Ancestral_sequence._data, subs_mat, -3, -.1, one_alignment_only=True)”* from Biopython and the coordinates of each ORF were transposed to correspond to the coordinates in the pairwise alignment. Subsequently, the Reading Frame Conservation^20,21^ (RFC) score was calculated for each ancestral ORF based on the pairwise alignment defined as: (length covered by the ancestral ORF aligned in the *S. cerevisiae* ORF frame)/(length of the *S. cerevisiae* ORF). For each ancestor of each ASR variation, we kept the ORF with the maximum RFC score.

To select the phylogenetic branch on which an ORF first appeared, based on a pre-defined RFC cut-off, we performed the following: Starting from the root of the phylogenetic tree and moving towards the leaves, we selected the first branch where an ancestral ORF existed with a maximum RFC higher than the pre-defined cut-off (e.g. 0.6). was selected as the evolutionary origin of the ORF. To select the phylogenetic branch on which an ORF first appeared, based on an empirical P-value, we performed the following: First, we computed an empirical P-value at each ancestral sequence of each ASR variation, by pseudo-randomizing the a given ancestral sequence 1000 times using the *random* package in Python 3 and then for each of the 1000 randomizations, ORFs (using the Stop-Stop definition only) were extracted, RFC was calculated for all, and the maximum RFC was kept, as described above. This resulted in a set of “randomized” 1000 best-RFC values, representing an empirical null model. Based on this distribution, we then calculated a P-value for the real maximum RFC score (one for each ancestral sequence of each ASR variation) by counting the number of randomized values greater than the real one and dividing by 1000.

### Statistical analyses

All statistics were done in R v3.6.2. Plots were generated using *ggplot2*^40^. All statistical details including the type of statistical test performed and exact value of n (n represents either number of genomes or number of genes) can be found in the Results and figure legends. Boxplots show median (horizontal line inside the box), first and third quartiles of data (lower and upper hinges) and values no further or lower than 1.5*distance between the first and third quartiles (upper and lower whisker). No methods were used to determine whether the data met assumptions of the statistical approaches.

## Data availability

The study uses publicly available data. Data source tables not included as supplementary information, as well as some scripts are available ***(will be once the work is accepted for publication)*** at https://github.com/Nikos22/

## Acknowledgements

We thank Saurin Parikh, Aaron Wacholder and all other members of the Carvunis lab for fruitful discussions that enriched this manuscript. The research project was supported by the Hellenic Foundation for Research and Innovation (H.F.R.I.) under the “3rd Call for H.F.R.I. Research Projects to support Post-Doctoral Researchers” to N.V. (Project Number:7330).

## Author information

### Contributions

ARC and NV conceived the study. ARC and NV wrote the manuscript. ARC supervised the study. NV, OA and VC performed the analyses.

### Affiliations

**Anne-Ruxandra Carvunis, Omer Acar, Vijay Cherupally**

Pittsburgh Center for Evolutionary Biology and Medicine, Department of Computational and Systems Biology, School of Medicine, University of Pittsburgh, Pittsburgh, PA 15213, United States.

**Nikolaos Vakirlis**

Institute For Fundamental Biomedical Research, B.S.R.C. “Alexander Fleming”, Vari, Greece

## Supplementary Figures

**Supplementary Figure 1:**
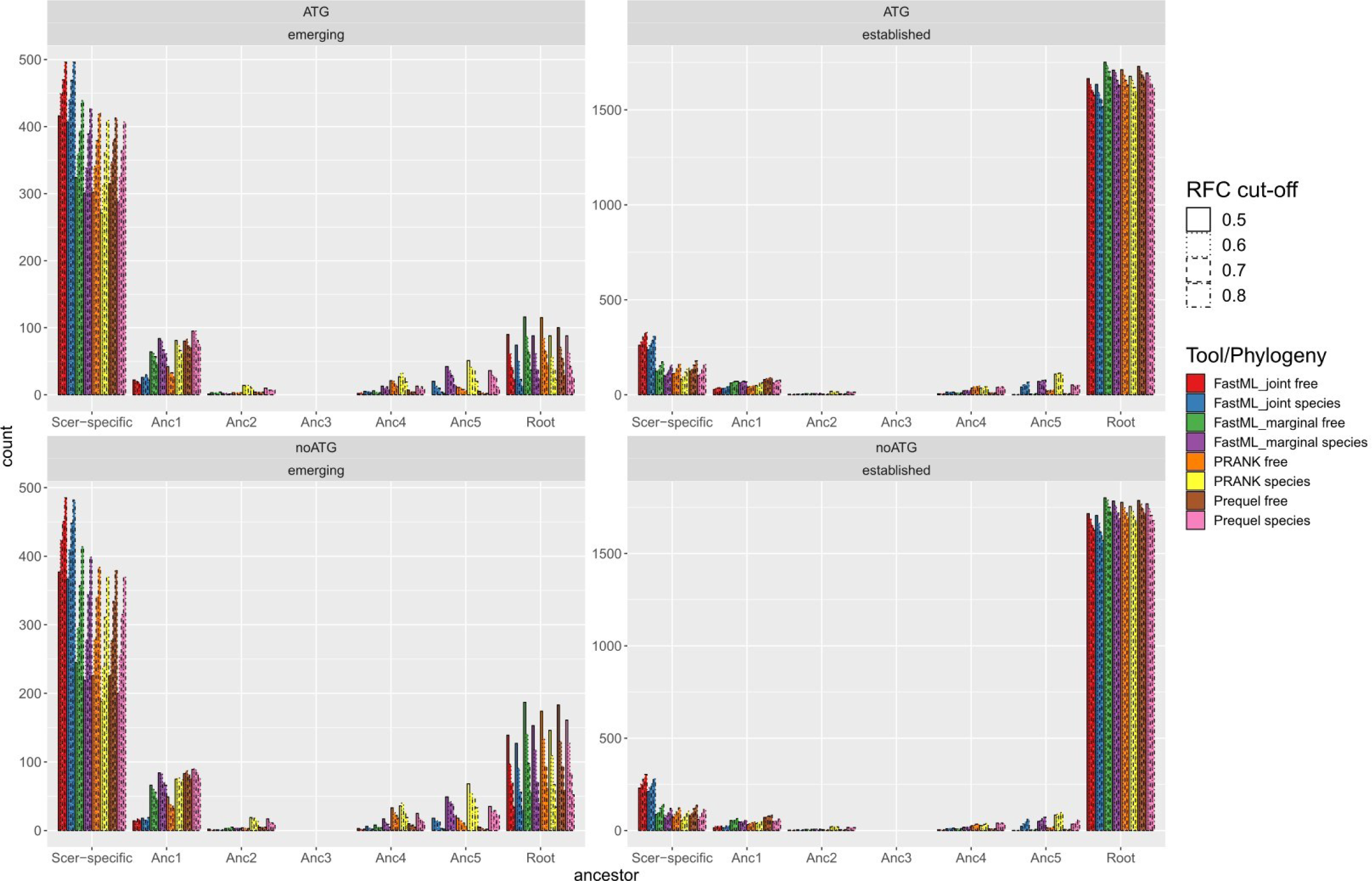
Distributions of branch where the most ancient ancestral ORF has been identified using four different RFC cut-offs and eight different ASR methodologies (tools+phylogeny). Bars within each node (e.g. “Root”) correspond, from left to right, to RFC cut-off of 0.5, 0.6, 0.7 and 0.8. Results for extended alignments using both “noATG” and “ATG” definition of ORFs are shown.

**Supplementary Figure 2:**
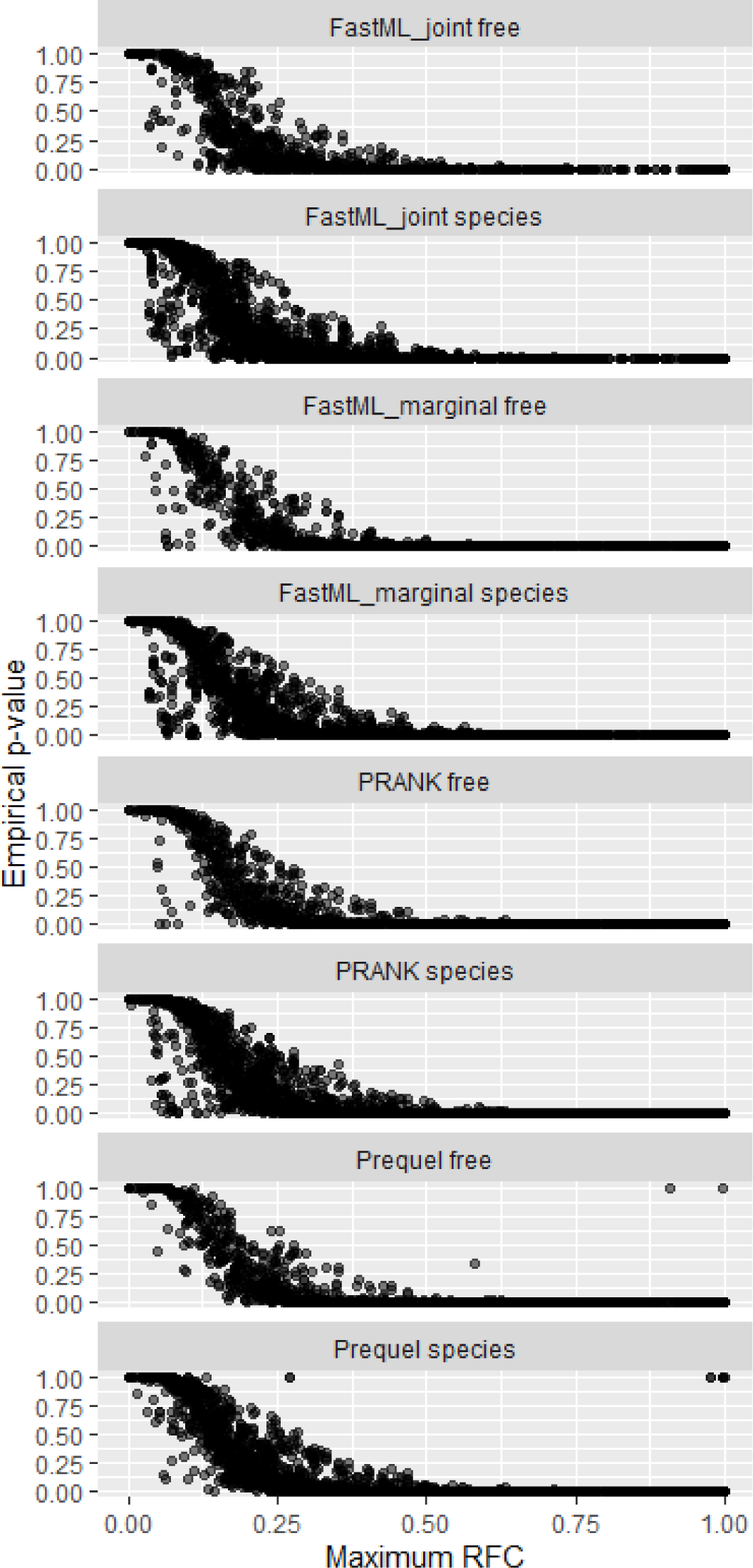
Relationship of the maximum (best) RFC score in the root ancestor and the associated empirical P-value for different methodologies, using the ORF-only alignments and the “noATG” ORF definition.

## Supplementary Tables

**Supplementary Table 1.** Gene names and associated data for 1,355 robustly ancient ORFs.

**Supplementary Table 2.** Best RFC and associated P-values for every ancestor, for the different methodological variations for 1,275 ORFs.

## Notes

### Competing Interest Statement

The authors have declared no competing interest.

